# Extreme Sampling for Genetic Rare Variant Association Analysis of Dichotomous Traits with Focus on Infectious Disease Susceptibility

**DOI:** 10.1101/2021.12.02.470949

**Authors:** MJ. Emond, T. Eoin West

## Abstract

As genomic sequencing becomes more accurate and less costly, large cohorts and consortiums of cohorts are providing high power for rare variant association studies for many conditions. When large sample sizes are not attainable and the phenotype under study is continuous, an extreme phenotypes design can provide high statistical power with a small to moderate sample size. We extend the extreme phenotypes design to the dichotomous infectious disease outcome by sampling on extremes of the pathogenic exposure instead of sampling on extremes of phenotype. We use a likelihood ratio test (LRT) to test the significance of association between infection status and presence of susceptibility rare variants. More than 10 billion simulations are studied to assess the method. The method results in high sample enrichment for rare variants affecting susceptibility. Greater than 90% power to detect rare variant associations is attained in reasonable scenarios. The ordinary case-control design requires orders of magnitude more samples to achieve the same power. The Type I error rate of the LRT is accurate even for p-values < 10^-7^. We find that erroroneous exposure assessment can lead to power loss more severe than excluding the observations with errors. Nevertheless, careful sampling on exposure extremes can make a study feasible by providing adequate statistical power. Limitations of this method are not unique to this design, and the power is never less than that of the ordinary case-control design. The method applies without modification to other dichotomous outcomes that have strong association with a continuous covariate.

## Introduction

As high throughput genomic sequencing has become available on a large scale, virtual armies of researchers and huge numbers of study participants have been contributing to the discovery of thousands of genetic associations, largely in common, chronic diseases among populations of European ancestry (1) (2). This is encouraging for progress in these areas of medicine and public health. It also has provided a wealth of knowledge on the technology and spurred development of statistical methods to handle large sample numbers from multiple ancestries. However, most diseases in the world remain untested for genomic associations, and most of these will not have the personnel and funding to assess more than a few hundred or even a few dozen genomes. Smaller sample sizes translate to less statistical power, especially for testing associations between phenotype and rare variants. Extreme phenotypes (EPs) designs have been put forth in the past as a means for providing the greatest statistical power under a fixed sample size(3) (4), and as few as 100 participants can give meaningful results. The “classical” EP design samples individuals from the extremes of the study trait in order to capture causal rare variants, enriching the sample with their presence relative to random sampling. At this time when interest in rare variants is swelling and small samples are getting less attention, we revisit the use of extreme sampling with attention to rare variants and infectious disease (ID). We review some results on classical use of EPs that utilizes sampling on extremes of phenotype. We then show how sampling on extremes of exposure rather than phenotype extends the principle of rare variant enrichment to infectious diseases. We then provide extensive simulation studies that show when the extreme exposure sampling is effective, emphasizing graphical displays to enhance communication of principles and results.

### The Essence of the Classical Extreme Phenotypes Design

In the context of genetic association studies, given a population of interest, the classical EP design samples from individuals exhibiting extreme values of the trait of interest. The underlying assumption here is that these extreme individuals are more likely to be carriers of variants that cause the extreme. Limiting sampling to the extremes limits the cost. Before discussing extensions of the EP design to infectious disease studies, it is enlightening to view an example where the classical trait-based EP sampling for a continuous outcome could have been used. The essence and potential effectiveness of the EP strategy are illustrated superbly by looking back on the findings for variants driving high plasma levels of low-density-lipoprotein cholesterol (LDL-C). Huijgen *et al* show that the population distribution of LDL-C is formed by a mixture of bell-shaped distributions corresponding to carriers of different LDL-C associated mutations(5). For the sake of illustration, we reproduce their findings for non-carriers and carriers of LDL-receptor (LDL-R) class I mutations. Fig. 1A shows histograms of these two sub-populations: two overlapping bell-shaped curves with different means and far fewer individuals in the more extreme sub-population. Fig. 1B shows the distributions combined, before genotype is known, conferring a longish right tail. In this example, random sampling from the population here would select mostly individuals from the middle of the distribution where most of the weight lies, resulting in 1 out of 9 individuals being carriers. In contrast, sampling individuals from the extreme, say individuals with LDL-C > 5mmol/L, increases that ratio by a factor of 80: 90% of those sampled from LDL-C > 5 will be carriers (Fig. 1A). A variant that is uncommon in the population becomes frequent in this EP sample. When those with LDL-C >5 mmol/L (“cases”) are compared to individuals with LDL-C < 2mmol/L (“controls”) in the opposite extreme of the distribution, Fisher’s exact test produces p < 1×10^-16^ for association of LDL-C with LDL-R class 1 mutations using just 50 high LDL-C cases and 50 low LDL-C controls. The power exceeds 99% here with 100 total participants. We use the term “enrichment” for the increase in variant frequency in the extremes.

The trade-off for this enrichment is that the odds ratio (OR) from the hypothetical case-control study above is biased relative to the population OR. However, that is of little concern when searching for associations that are difficult to detect for variants that will inevitably undergo functional testing and estimation of prevalence in various populations. Our focus here is on power to find causal genetic variants and not on precise estimation of their population effect. Note that the significance test (e.g. Fisher’s exact test above) is unbiased, meaning that when no association exists, the p-value will not show spurious significance from the biased sampling. Fig. 1 helps illustrate this: if there were no mutations that drive up LDL-C in some people, the red distribution would not exist, and sampling from the extremes would result in sampling all variants from the blue distribution. The result is a null comparison as there is no difference in variant frequency between the individuals in the two tails of blue distribution.

**Fig 1.**
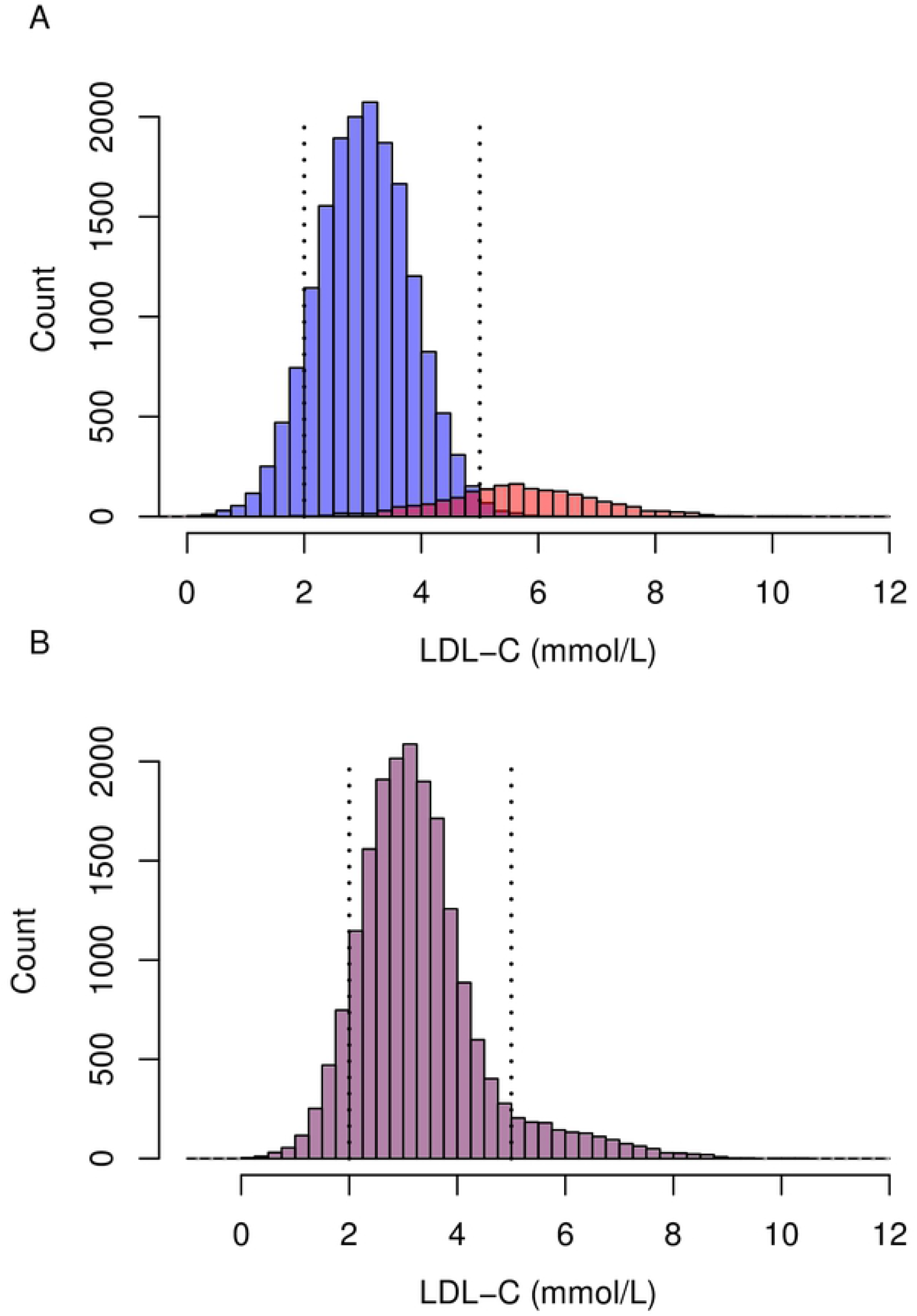
Extreme Phenotypes for a Continuous Trait. (A) The histograms of low-density lipoprotein cholesterol (LDL-C) for people with and without LDL-R type I mutations are overlaid. This illustrates well how sampling the extremes of LDL-C results in a higher proportion of rare variants selected into the sample compared to ordinary case-control sampling. Red=LDL-R variant carriers, blue = no LDL-R variants. From Huijgen (11) et al. (B) The population distribution of LDL-C is a mixture of distributions for carriers and non-carriers.

### Extreme Sampling for Infectious Disease Association Studies

The most germane and biologically natural outcome for an the study of an infectious disease is the dichotomous state of infected or not infected, and it is of interest to find genetic variants that protect one from infection or make one much more susceptible. While the dichotomous outcome defines the case and control groups (infected and not infected), it usually cannot be further dissected into extremes. On the other hand, the exposure to the infectious agent is in theory a continuous (or semi-quantitative) variable that can be used to define extreme groups: cases (infected) with low exposure and controls (uninfected) with high exposure. The group status is then tested for association with genotype. Historically, the discovery of CCR5Δ32 as a protective variant for HIV-1 infection is a demonstration of this design by happenstance(6) (7) (8).

Researchers noticed some hemophiliacs remained uninfected for HIV-1 even after multiple infusions with virus-contaminated blood products. These individuals constitute a resistant extreme. In this example, the exposure to HIV-1 was well-documented and quantifiable. In a second example of high interest, the sera of a group of 12 children who were observed to be resistant to severe malaria were compared to that from 11 children who were susceptible (from a total cohort of 784 children)(9). Intense quantitative analysis led to discovery of antibodies that prevent the malaria parasite, *Plasmodium falciparum,* from reproducing in the hosts’ blood, providing a protective effect. While the latter is not strictly genomic, the design principle is completely parallel with that for extreme exposure genetic association testing. Both examples illustrate the role of astute clinical and field observation along with the value of extreme sampling. Note that the term “exposure” throughout this paper refers to pathogen exposure and is not to be confused with genotype as the exposure in a genetic association test. We use the terms “genotype” or “carrier/non-carrier” to denote genetic status. With these examples in mind, we formally extend the extreme sampling for genetic association studies for susceptibility to infectious diseases in general.

## Methods

### Overview

Execution of extreme exposure sampling is simple in theory. Cases (infected individuals) with low exposure are selected for study along with controls (uninfected) individuals with high exposure. Logistic regression is then applied using infection status as the outcome and genetic score as the independent variable of interest (the genetic score for each locus). We expect the cases to be enriched for deleterious/disease-causing variants and expect the controls to be enriched for protective variants. The likelihood ratio test (LRT) for logistic regression is used to test for association between disease and genetic score. We use simulations to illustrate the principle of enrichment via sampling on pathogen exposure level and to assess the behavior of the method. The “extremeness” of the exposure is measured by percentiles of the exposure distribution.

### Simulated Data Generation

Infection status is determined by the level of pathogen exposure in combination with genotype for causal and protective variants. For these studies, to provide realistic results, we generated data to approximate the exposure/genotype/outcome relationship that we observed in a study of HIV-1 susceptibility. More specificially, let H denote the exposure level for an individual and let G denote presence of a variant at the locus being tested (G=0 or 1). We take the state of nature to be such that the probability of being infected, denoted by μ is dependent on H through the commonly used liner logistic function when G=0: logit(μ_0_) = −30 + H, or equivalently, μ_0_=exp(α+H)/(1+exp(α+H)). The population relative risk (RR) for carriers versus non-carriers is μ_1_ /μ_0_ where the subscripts denote presence (1) and absence of a susceptibility variant (0). Hence, the dependent variable, Y (=1 for cases, =0 for controls) has mean (α+H)/(1+exp(α+H)) when G=0 and mean RR x (α+H)/(1+exp(α+H)) when G=1. We generate H and G independently then generate a population of Y’s, each as a random binomial variable with the foregoing mean. We take H to be normally distributed with mean 20 and standard deviation (SD) 10; units for H do not affect the results, but values of H that are less than zero are omitted. G is generated for each individual according to the minor allele frequency (MAF) to be studied, and then Y is generated from G, H and the RR to be studied. We vary the MAF and RR across simulations as well as the extremeness of sampling as measured by the percentiles of H. The model for H is held fixed for comparisons. This roughly replicates a model for the probability of seroconverting to HIV-1 positive within 1 year given H is the number of unprotected sexual encounters with an HIV-1 infected partner(10) for an average at-risk individual. We give examples using MAF=0.005 and MAF=0.001, but it is critical to note that these can be the cumulative MAF of several variants that are counted as one genetic unit in order to increase the frequency (or “burden”), as originally conceived by Morris and Zeggini (11) and as in applied in Emond et al (12). Without loss of generality, we assume a dominant model. The complete step-by-step algorithm for generation of the simulation data is provided as Supplementary item **S1**.

### Testing for Association

Controls are sampled from highly exposed individuals who remained uninfected while cases are sampled from infected individuals with low exposure. Tests of association between genotype and outcome are applied and results tallied. For this study, unadjusted logistic regression using the LRT with G as the single covariate is performed along with Fisher’s exact test and Firth regression. All test results are for a Type I error rate of 5×10^-8^ Firth regression is chosen because of the small numbers of variant carriers in some instances, and Firth regression is purportedly more stable in these instances(13) (14). The LRT was suggested by Morris and Zeggini, and we have found this test to be very reliable with small samples(11). Random samples of cases and controls are assessed for each RR+MAF combination for comparison to the results for extreme sampling on exposure. Test size was evaluated by performing 10 billion LRT tests under the null hypothesis with only 50 cases and 50 controls. To test the robustness of the results to errors in the exposure measurement, we use the scenario of MAF=0.005, N=300 per group and sampling from the top and bottom 1%. To simulate types of errors we have observed, we insert a portion of samples with random exposure rather than extreme exposure, and examine a range of sample portions with this random exposure. Except where noted, all simulation results are the mean of 500 repetitions, drawn from a simulated population of 1.2 million. Percentiles refer to population percentiles and not sample percentiles.

To illustrate a scenario in which very high power is simply not attainable, we use a population size of 50,000 in a series of simulations with MAF=0.005 and RR=0.5.

## Results

We now have rare variant association results from simulated data where the outcome depends on both pathogen exposure and genotype; we have sampled cases with low exposure and controls with high exposure and tested for association of outcome with genotype. One of the results of the sampling is that individuals with high pathogen exposure who would be destined to become infected in the absence of the protective variant are much more likely to be controls when the variant is present (G=1). This produces a tail on the distribution of H among controls, with the size of the tail depending on the MAF of G, the distribution of H and the population relative risk (RR). This tail phenomenon, similar to Fig. 1 but mathematically different, is illustrated in Fig. 2 for the model above and several values of the population RR. While in Fig. 2A we see that variant carriers are a miniscule portion of the uninfected controls, as expected for MAF=0.005, Figs. 2B and 2C show that it is possible to sample from Hs that are so extreme (above 40, say) that all of the sampled individuals will be carriers of the protective variant (G=1). Fig. 2D shows the distributions of the infected and uninfected carriers and non-carriers on the density scale for comparison. As expected, overall, uninfected individuals (red) have a larger probability of having small values of H compared to the infected (blue). The controls have distinctly different distributions for carriers and non-carriers (red vs aqua) because G is protective, while the carrier and non-carrier distributions among cases have no perceptible difference (which would not be true for a susceptibility variant.)

**Fig 2.**
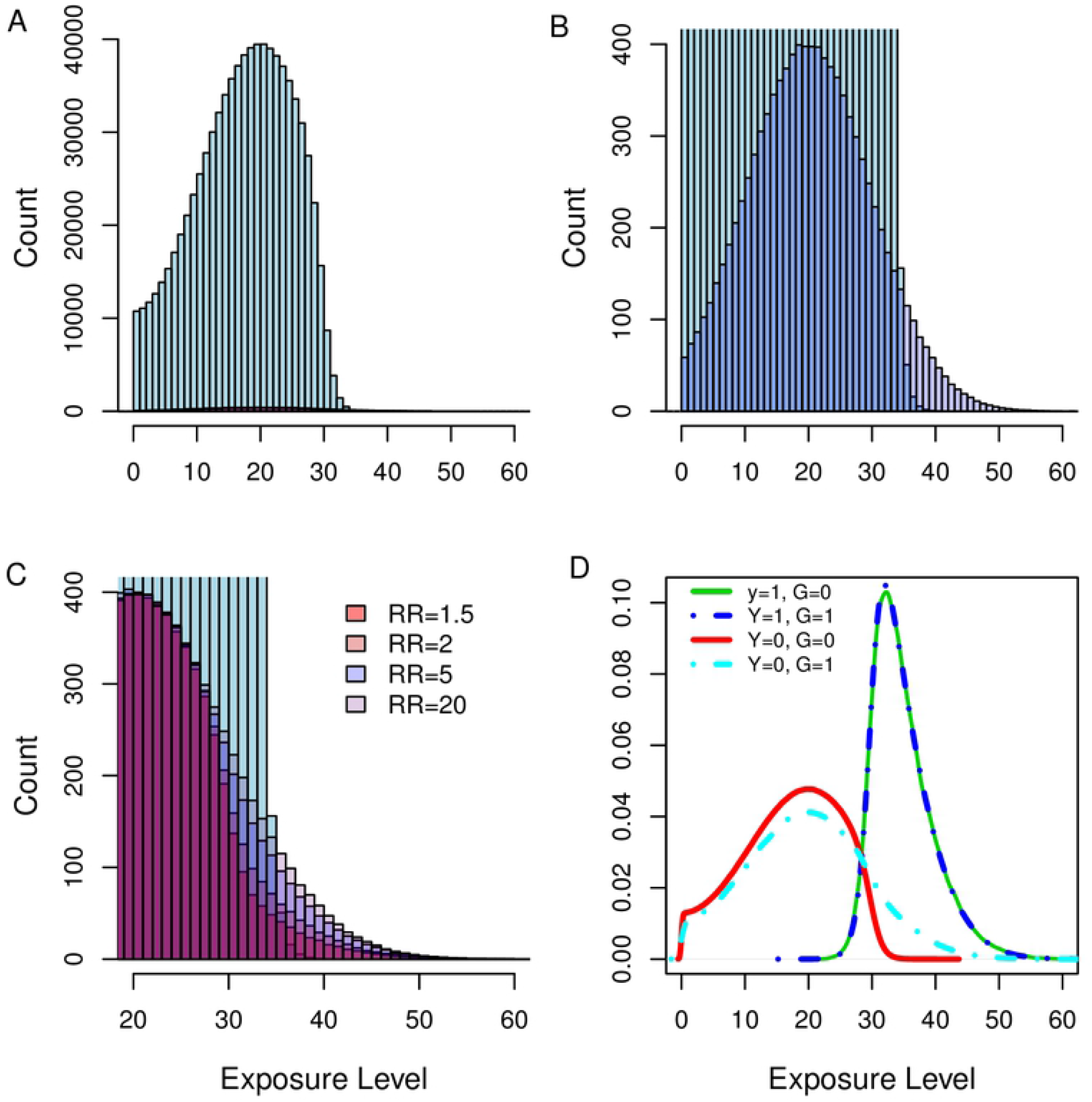
Population Distributions of Exposure Level, H, for Different Groups. (A) Histograms of H for individuals with Y=0 and G=0 (blue) and for those with Y=0 and G=1 (red, barely discernable). The minor allele frequency for the protective variant is 0.005 in all panels. (B) Zoom of Fig 2A. showing the carrier distribution (red) and non-carrier distribution (blue) for the protective variant. Sampling above an exposure value of 40 (2 SD above the mean) would result in virtually all controls being carriers. (C) Further zoom of Y=0, G=0 (light blue) and Y=0, G=1 (red shades) for four values of the population relative risk for four different variants. RR=1.5, 2, 5 and 20 (darkest to lightest red). The plot shows little gain in enrichment capacity after RR=5. (D) Distribution of H on the density scale. All four carrier-by-outcome groups are shown for RR=20. Y=1, G=0 (solid green); Y=1, G=1 (dotted blue); Y=0, G=0 (red); Y=0, G=1 (dotted aqua). The heavy tail for Y=1, G=1 allows enrichment of the sample by selecting controls with high values of H.

Table 1 shows sample size results for MAF=0.001 and 0.005, RR’s of 0.167, 0.25 and 0.5, and percentiles of H ranging from the 0.999 to 0.95 on the high side (0.001 to 0.05 on the low side). Sample sizes are given for four levels of power at a Type I error rate of 5×10^-8^. Additional power results are given in S2 Tables. Power increases as the RR deviates further from one, so results for more extreme RR’s can be inferred from Table 1. A RR of 1/6 (0.167) is a reasonable effect size for an RV in an ID. Sampling within the 95th and 5th percentiles attains near 100% power to detect a RR=0.167 with 1394 per group when the MAF=0.005 or greater (Table 1, row 7). On the other hand, random sampling would require > 5000 per group to achieve just 80% power, a huge difference. In general, sample size needs decrease as the sampling percentile and/or the RR become more extreme and as the MAF becomes less extreme (Table 1). Table 1 also highlights the potentially exquisite power of this method, with fewer than 100 individuals per group needed for 95% power to detect RRs at 0.5 or less when sampling below the 0.001 percentile and above the 0.999 percentile. This is in stark contrast to the >7000 needed for random cases and controls (Table 1; S2). The empirical OR in column 9 is the OR observed within the extreme sample of the simulation (the mean of 500 trials). This is another measure of how successful the enrichment sampling is: for row one in Table 1, the empirical OR=0.001, meaning that a case is about 1/1000 as likely as a control to be a protective rare variant carrier in the extreme sample, compared to 1/6 (RR=1/6) as likely in the general population. Some of the combinations in Table 1 are quite extreme, but a very reasonable scenario is one in which RR ~ 0.25, MAF≥0.005 and samples in the top and bottom 1% are available (Table 1, row); power is 80% with 287 individuals per group.

**Table 1.**
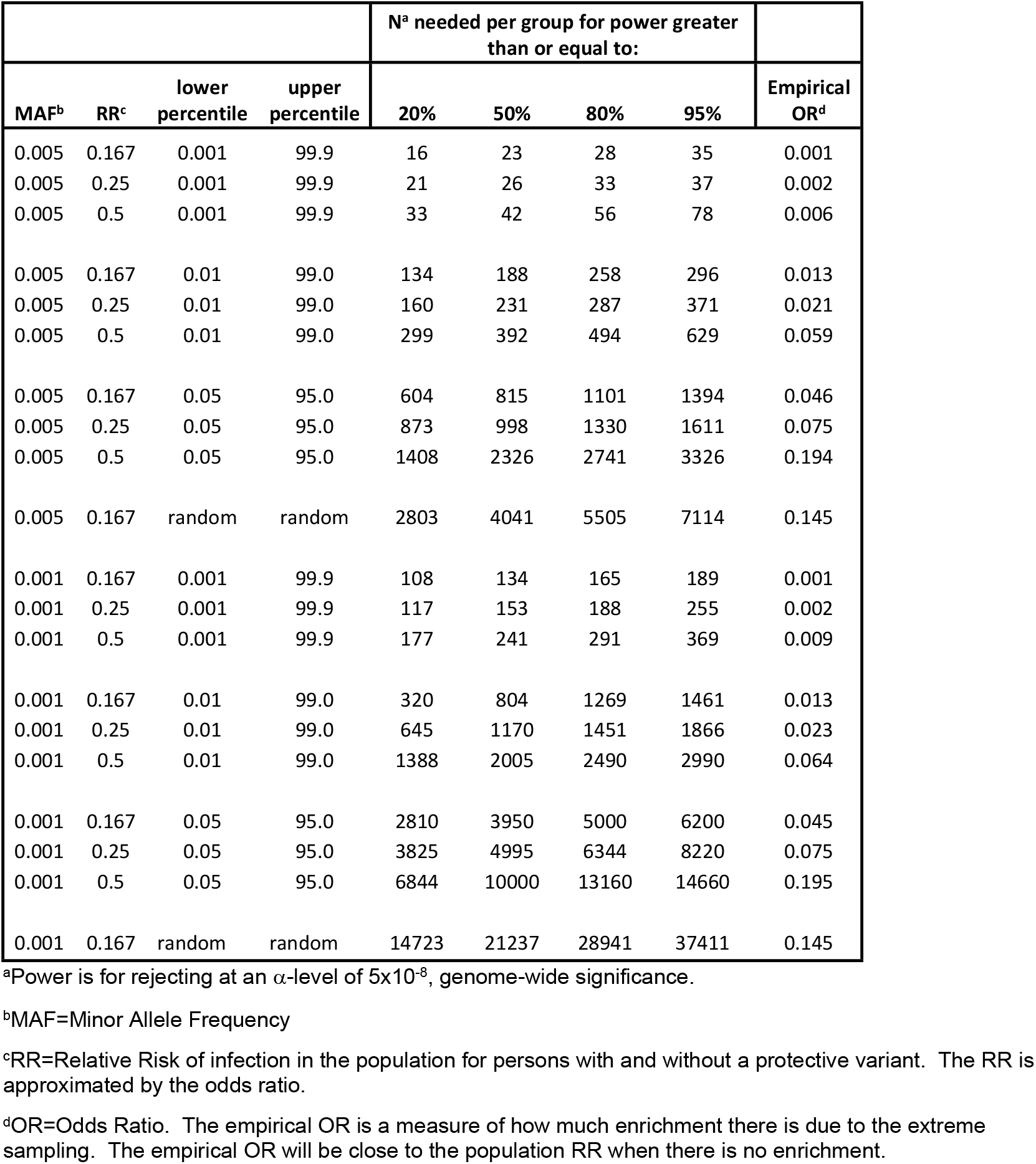
Sample sizes Needed for Case and Control Groups to Provide the Indicated Power Under the Extreme Sampling Scenario in Columns 3 and 4.

As expected, Fisher’s exact test is less powerful than logistic regression with the likelihood ratio test; but, unexpectedly, Firth regression provided no benefit over ordinary logistic regression and was consistently less powerful the the LRT (S2 Table).

Test sizes from 0.05 to 5×10^-8^ were evaluated by performing 10^10^ LRT tests under the null and then tabulating how many results showed p<α for α=0.05, α=0.005,…, α=5×10^-8^. Comparing observed and expected, the LRT test was slightly optimistic at higher p-values but was conservative for genome-wide significance levels (Fig. 3).

**Fig 3.**
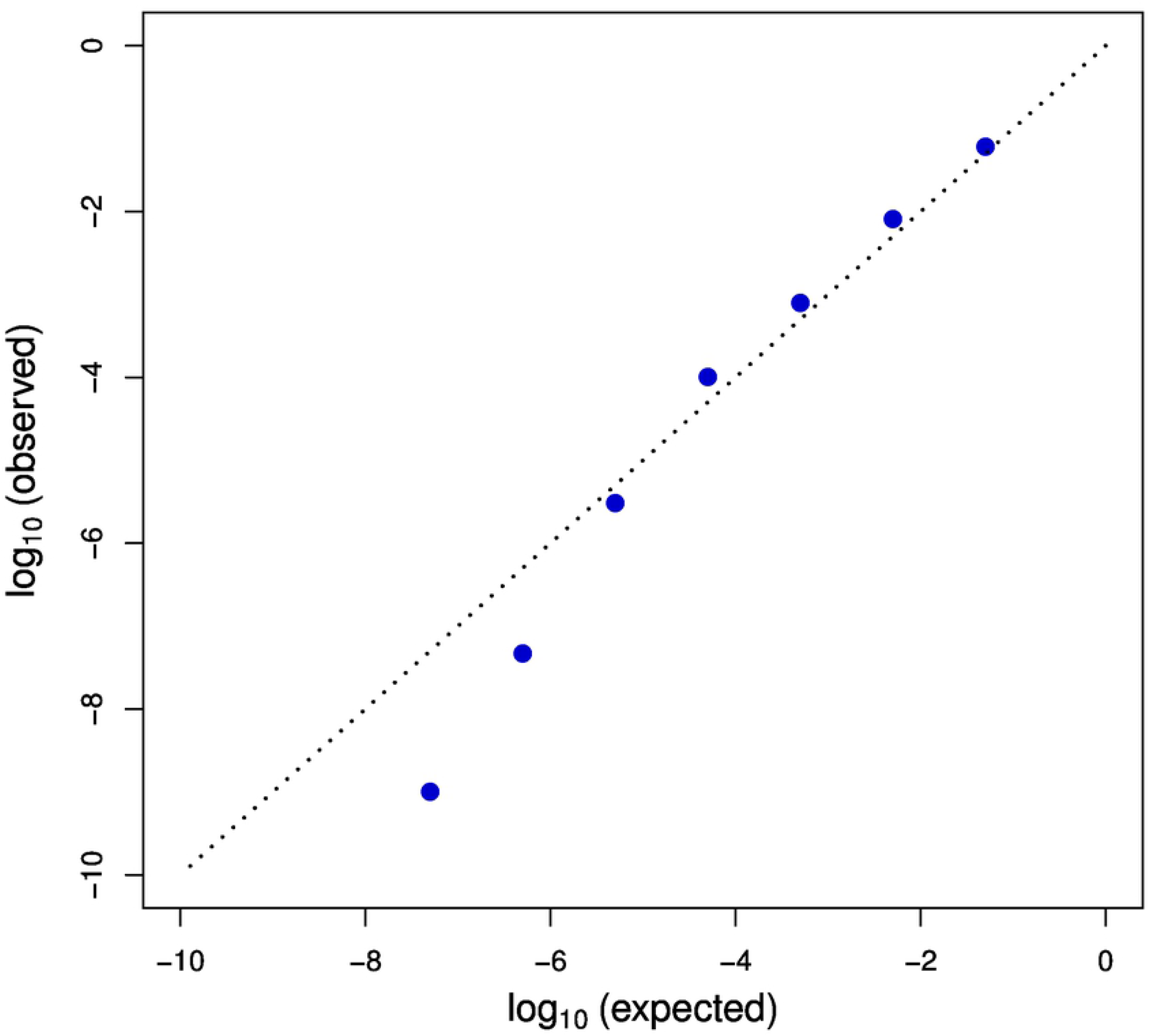
Observed vs. Theoretical Test Sizes for the Likelihood Ratio Test. For protective variants, it is known that a case:control ratio larger than 1 can provide additional power for logistic regression with the Wald test under random sampling(15). To see whether this holds true for extreme sampling with logistic regression and the LRT, we performed a large simulation study in which we varied the case:control ratio while keeping the sample size fixed. The optimal case:control ratio is approximately 1/OR^1/2^ for logistic regression using the Wald test when cases and controls are sampled randomly(15). We found that the optimal case:control ratio was 4:3 for two different values of the RR (0.5 and 0.11, resulting in predicted maximums at 1.4 and 3.0) and two different overall sample sizes (Fig. 4) for both logistic regression and Fisher’s exact test. The case:control ratio providing maximal power was stable over the range of scenarios. The result was the same for RR = 0.04, an extreme value that should show an effect on the optimal case:control ratio if there was one (results not shown).

In a previous study of genetic risk factors for HIV-1 infection, we noticed discrepancies in reports of exposure to unprotected sex when reports were obtained from the different partners(10), along with outcomes that were inconsistent with some recorded exposures. Sampling based on these erroneous exposure reports would put non-extreme individuals in place of extreme individuals. To investigate the effect of this, we used 300 individuals per group with sampling at the 99^th^ percentile and then substituted random samples for extreme samples in increasingly large numbers. Fig. 5A shows the effect on power is quite deleterious, with power decreasing to *α* as the number of observations with error nears 50% of the sample size. In fact, power is slightly better, unexpectedly, when the samples with error are discarded (Fig. 4B).

**Fig 4.**
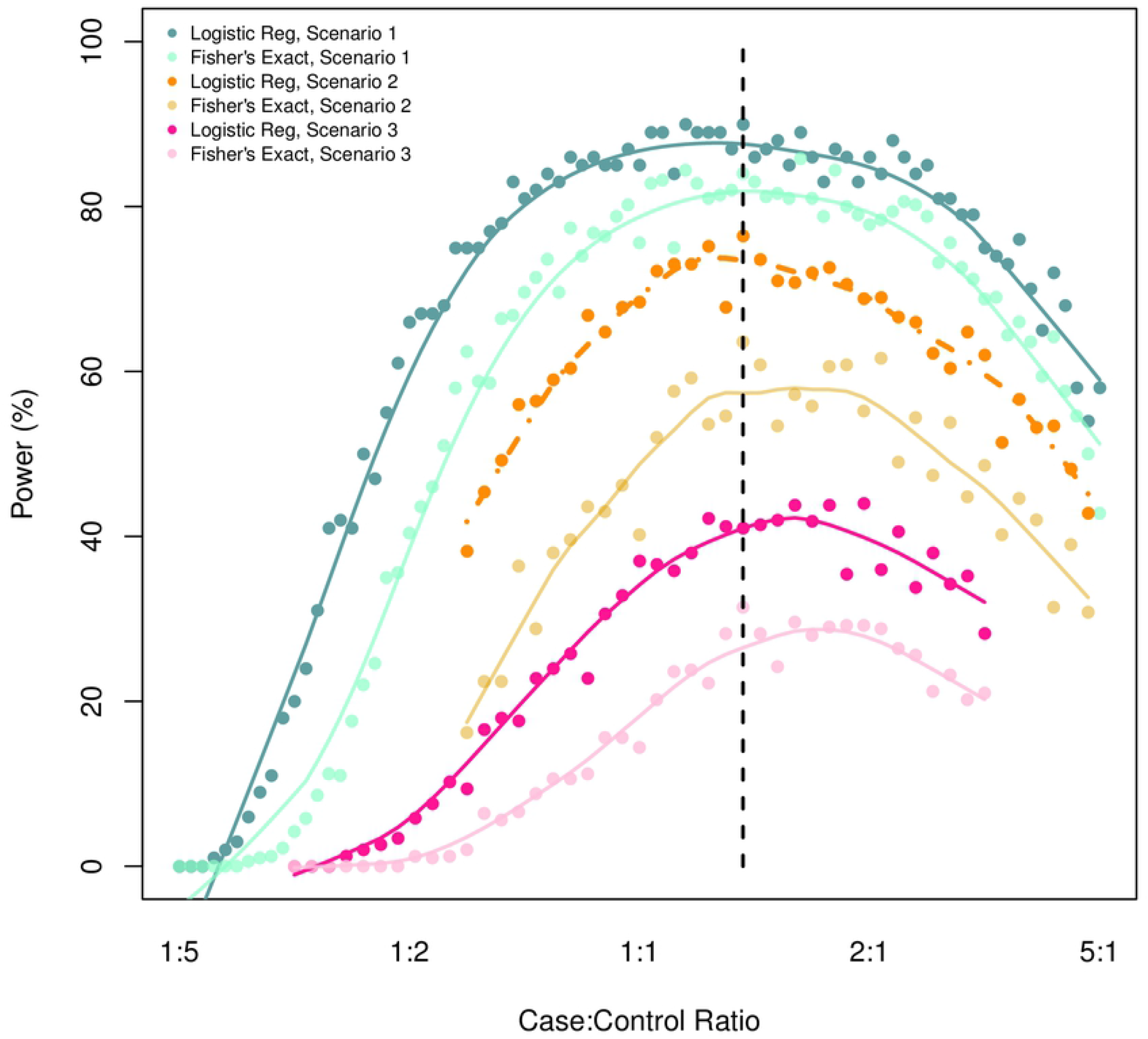
Power vs Case:Control Ratio for a fixed sample size. The best power occurs when samples are spread more or less evenly among cases and controls. No appreciable power gain is found for allotting more samples to controls as would be true for ordinary logistic regression and the Wald test. Scenario 1 (blues): RR=0.5, N=600; scenario 2 (oranges): RR=0.25, N=400; scenario 3 (reds): RR=1/9, N=400; all were sampled at the 1^st^ and 99^th^ percentiles.

**Fig 5.**
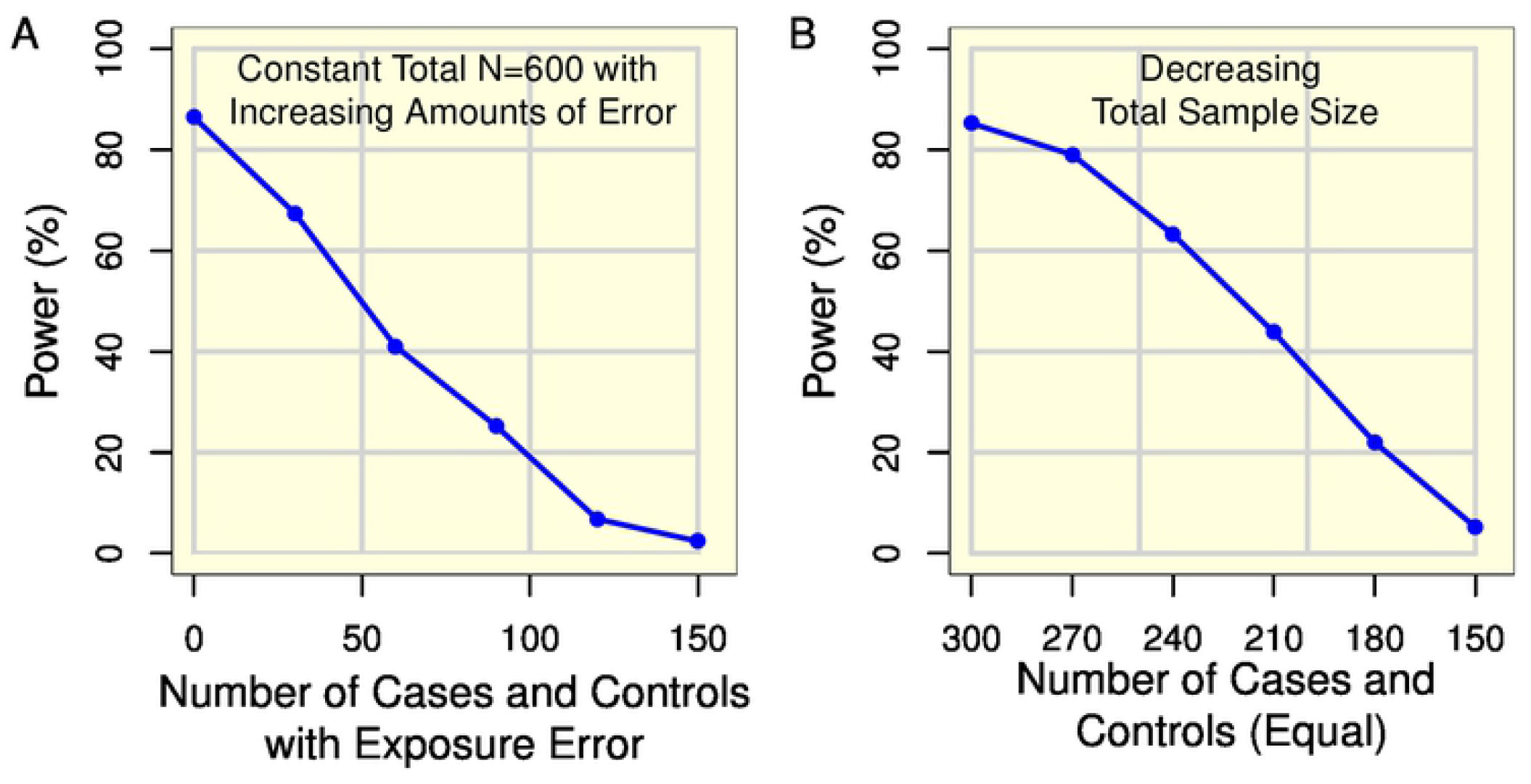
Effect of Mismeasuring Extreme Exposure. (A) Starting with 300 extreme exposure individuals per group, portions of each group were replaced by randomly chosen cases and controls (with their corresponding values of G.) Power decreases drastically with increasing portions of mismeasured (non-extreme) samples. (B) Power for the same scenario as in A. except observations with mismeasured values are discarded. Power is better than when including the randomly mismeasured samples.

It is especially important to consider the underlying population in an extreme sampling study because if the size of the population is small, this can limit the extremes. Table 2 shows the results for a population of size 50,000 when trying to detect a variant or variants with cumulative MAF=0.005. The maximum attainable power here is only 77.8% with n=400 per group; further increases in sample size actually result in decreased power, an especially noteworthy result. The empirical OR increases with every increase in sample size here, becoming less and less extreme because more extreme samples don’t exist.

**Table 2.** Aberrant Power Behavior.

| MAF | RR | %ile | N cases | N Controls | Power (%) |  |  |  |
| --- | --- | --- | --- | --- | --- | --- | --- | --- |
|  |  |  |  |  | Fisher's Exact Test | Logistic Regression | Firth Regression | Empirical OR |
| 0.005 | 0.5 | 99.96 | 20 | 20 | 28.6 | 55.0 | 41.6 | 0.001 |
| 0.005 | 0.5 | 99.94 | 30 | 30 | 40.6 | 70.4 | 51.2 | 0.003 |
| 0.005 | 0.5 | 99.92 | 40 | 40 | 44.4 | 65.2 | 54.8 | 0.005 |
| 0.005 | 0.5 | 99.90 | 50 | 50 | 51.8 | 70.0 | 61.2 | 0.006 |
| 0.005 | 0.5 | 99.80 | 100 | 100 | 48.6 | 69.8 | 60.4 | 0.014 |
| 0.005 | 0.5 | 99.60 | 200 | 200 | 61.8 | 75.4 | 69.2 | 0.027 |
| 0.005 | 0.5 | 99.40 | 300 | 300 | 65.8 | 75.4 | 70.8 | 0.041 |
| 0.005 | 0.5 | 99.20 | 400 | 400 | 66.8 | 77.8 | 74.6 | 0.049 |
| 0.005 | 0.5 | 99.00 | 500 | 500 | 67.0 | 77.6 | 73.6 | 0.060 |
| 0.005 | 0.5 | 98.00 | 1000 | 1000 | 69.4 | 75.4 | 71.8 | 0.105 |
| 0.005 | 0.5 | 96.00 | 2000 | 2000 | 65.4 | 73.6 | 71.2 | 0.168 |
| 0.005 | 0.5 | 94.00 | 3000 | 3000 | 64.4 | 71.0 | 69.0 | 0.214 |
| 0.005 | 0.5 | 92.00 | 4000 | 4000 | 63.4 | 69.0 | 67.8 | 0.256 |
| 0.005 | 0.5 | 90.00 | 5000 | 5000 | 65.0 | 69.4 | 68.8 | 0.282 |
| 0.005 | 0.5 | 60.00 | 20000 | 20000 | 44.4 | 46.8 | 44.8 | 0.421 |

## Discussion

We have shown here that, by sampling on extremes of pathogen exposure, very high statistical power can be attained for testing associations of rare variants with infection status. While sampling is necessarily quite extreme in some cases, such extremes can be obtained(16). In addition, RVs are more likely to be functional, which increases the motivation to enrich the sample with RVs by extreme sampling(16).

Some of the sample sizes in Tables 1 and S2 are in the thousands and perhaps in a cost region above that of many researchers, but random sampling, as expected, does no better: with 5000 samples per group needed for 77% power with MAF = 0.005 for detecting an RR=0.17 (S1A Table, Scenario L), and with 20,000 per group, the power is only 51% for MAF=0.001 and RR=0.17 (S1B Table, Scenario Y). Extreme sampling might be the only hope of attaining good power. Power of 80% or higher is advisable for a replication study, but for a first stage agnostic, genome-wide or exome-wide study, lower power thresholds are still quite useful when several variants are expected to play a role. In the latter situation (which is believed true for most IDs(17) (2)), the multiplicity of variants provides multiple chances for uncovering a causal variant at a genome-wide significance level and a low chance of missing all of them. With a power of 20% for each of 12 variants, for example, the probability of finding at least one of them is 1-(1-0.2)^10^ = 0.93. In fact, we advocate the use the False Discovery Rate (FDR)(18) rather than p-values in phase I discovery studies along with filtering variants for functionality, discarding before analysis variants that are unlikely to have functional role in the infection. For example, synonymous variants could be discarded. These two measures further increase power to identify RV associations. Because the FDR is a one-to-one function of the p-value, the general principles found here hold for the FDR. Power calculators for case-control studies can be used to estimate the power of the extreme sampling design if the empirical odds ratio (Table 1) can be estimated.

Table 1 might not be strictly applicable to others’ research scenarios but serves to illustrate a set of scenarios, both attainable and perhaps not attainable. It is important to understand the latter as much as the former. High power is attained at the cost of careful sampling and searching in the population to find the extremes. Fig. 3, degradation of power as the error rate increases, emphasizes that substituting individuals with random exposures or unknown exposures should be avoided in the extreme design. Though it is not always possible to identify such errors, even suspicious samples should be avoided. While all measurements have some error, the type of error here is large in the sense that a random value of H from the population is used in place of an extreme value, resulting in relatively large deviation between the assume value of H and the actual value. The observed effect is consistent with the findings of Pelosos et al where they find that augmenting 200 extreme cases and 200 extreme controls with 1000 random samples provided little gain in power and resulted in power loss if the latter samples were not down-weighted(16). More study of different kinds of errors is warranted, along with development of validation sample methods to statistically correct errors(19).

The age of the individual at infection can be an important aspect of the exposure measurement. Such a situation includes *Pseudomonas aeruginosa* infection cystic fibrosis (CF) affected individuals(12)·(20). In CF, the exposure to *Pseudomonas* in the environment is more or less constant over time, so older children have greater exposure; remaining infection-free into one’s twenties is a rare (extreme) event under genetic influence(21) (22).

Focusing on a rarer subgroup of outcomes can also be helpful in achieving enrichment of the sample with RVs. The infection under study might have distinct, rare manifestations that are of clinical relevance and that arise only rarely in some infected individuals. Examples are neurological involvement in syphilis and West Nile virus, and development of certain post-acute symptoms following COVID-19 infection.

In order to have faith in our power estimates, the test size should be correct. That is, when the null hypothesis is true, the proportion of tests with p-value < α should be quite close to α. Inflated test sizes (anti-conservative results) are often seen with small samples and very low p-values. This is not a problem here: we have shown through 10 billion simulations that the test size for logistic regression using the LRT p-value has correct size with only 50 cases and 50 controls at p-values as low as 5×10^-8^. This is itself an important result.

The scenario for which the population size is only 50,000 and the power decreases after 400 per group is not just a toy example. Isolated populations with unique background genetics in combination with rarity of the sought-after variants will lead to such a scenarios in reality. This problem is not limited to extreme sampling.

One should not adjust for pathogen exposure when using this method. The method shown here is equivalent to performing logistic regression of disease status on H and then sampling the most extreme residuals as in Guey et al(23). The method in Guey et al might be used as an alternative to finding the extremes of exposure itself, but in all examples shown here the Guey method fails because of complete separation of cases and controls resulting in lack of convergence of the logistic regression model. Guey et al also use a significance level of 0.001, whereas the work here informs us on genome-wide significance levels, a critical addition.

One should adjust for other confounders when they exist. In particular, adjusting for ancestry should be done(24).

We found that the optimal case:control ratio was empirically 4:3 for different overall sample sizes, different tests and different values of the OR (0.5 and 0.11). The predicted optimal ratio under random sampling is 1/OR^1/2^, but our results do not reflect that prediction. Staying with a 1:1 case:control ratio seems advisable and provides the best strategy for finding both protective and deleterious variants.

We have purposely used a model with only one RV for simplicity. We chose to illustrate this method using a protective variant, but by symmetry of the logistic model, the general methods and results also apply to deleterious variants. Further, the method applies to other dichotomous outcomes that are strongly correlated with a continuous covariate.

A limitation of this method is that quantification of the infective agent often is not available. However, it might be possible to obtain quantified exposure measurements in situations where they are not routinely made. For example, aerosolized and windblown *Burkholderia pseudomallei,* an environmental saprophyte in certain tropical regions that causes the ID melioidosis, is correlated with melioidosis incidence(25). The melioidosis incidence is too low even in the aerosol-exposed areas for this metric alone to be useful in identifying particularly resistant individuals, but it *can* be useful in identifying highly susceptible individuals who could be compared with exposed, uninfected family members using a paired test. We emphasize that controls in any genetic susceptibility study must be exposed. In other important situations, extremes can be identified with non-continuous exposure information. These situations include certain infectious hemorrhagic fevers where just one exposure to bodily fluids or mucous membranes of an infected individual confers extremely high risk for infection, so that individuals who avoid infection after such exposure can be considered extreme.

Another limitation of this study is that the model used here for Y as a function of H (logistic model) might not be close enough to the situation under study for Table 1 to apply. However, the procedures used here can be repeated for the investigator’s model.

When sample sizes cannot be in the 10’s of thousands, careful observation and extreme sampling can improve power and require far fewer samples. We argue extreme sampling should be considered more frequently given its high power along with the growing ability to technically identify and genotype rare functional variants, the need for progress in resource-limited areas, and growing antibiotic resistance and emergence of serious new infections worldwide.

